# *CaProDH2*-mediated modulation of proline metabolism confers tolerance to Ascochyta in chickpea under drought

**DOI:** 10.1101/2021.06.25.449950

**Authors:** Mahesh Patil, Prachi Pandey, Vadivelmurugan Irrulappan, Anuradha Singh, Praveen Verma, Ashish Ranjan, Muthappa Senthil-Kumar

## Abstract

Drought and leaf blight caused by the fungus *Ascochyta rabiei* often co-occur in chickpea (*Cicer arietinum*)-producing areas. While the responses of chickpea to either drought or *A. rabiei* infection have been extensively studied, their combined effect on plant defense mechanisms is unknown. Fine modulation of stress-induced signaling pathways under combined stress is an important stress adaptation mechanism that warrants a better understanding. Here we show that drought facilitates resistance against *A. rabiei* infection in chickpea. The analysis of proline levels and gene expression profiling of its biosynthetic pathway under combined drought and *A. rabiei* infection revealed the gene encoding proline dehydrogenase (CaProDH2) as a strong candidate conferring resistance to *A. rabiei* infection. Transcript levels of *CaProDH2*, pyrroline-5-carboxylate (P5C) quantification, and measurement of mitochondrial reactive oxygen species (ROS) production showed that fine modulation of the proline–P5C cycle determines the observed resistance. In addition, *CaProDH2-*silenced plants lost basal resistance to *A. rabiei* infection induced by drought, while overexpression of the gene conferred higher resistance to the fungus. We suggest that the drought-induced accumulation of proline in the cytosol helps maintain cell turgor and raises mitochondrial P5C contents by a *CaProDH2*-mediated step, which results in ROS production that boosts plant defense responses and confers resistance to *A. rabiei* infection. Our findings indicate that manipulating the proline–P5C pathway may be a possible strategy for improving stress tolerance in plants suffering from combined drought and *A. rabiei* infection.

## Introduction

Drought exerts complex effects on plant diseases. The effects of drought on pathogen infection may be positive or negative. The factors governing the net outcome of combined stresses are the intensity of each imposed stress, the order in which they are imposed, and the pathogen type (Ramegowda and Senthil-Kumar, 2015; Pandey et al., 2017). Drought co-occurring with fungal infections affects both growth and productivity in chickpea (*Cicer arietinum*). While low soil moisture content increases the severity of black root rot (Bhatti and Kraft, 1992; Sharma and Pande, 2013; Sinha et al., 2019), it reduces collar rot incidence by inhibiting fungal colonization inside plants (Tarafdar et al., 2018). Drought stress also modulates the interaction between foliar pathogens and plants. For example, a survey of Ascochyta blight in 251 chickpea fields (including experimental and farmer fields) in Ethiopia indicated a low disease incidence during the dry year 2015–2016 (Tadesse et al., 2017).

Ascochyta blight, caused by *Ascochyta rabiei*, may cause complete harvest loss under cold and humid weather conditions (Pande et al., 2005; Jaiswal et al., 2012; Sharma and Ghosh, 2016). *A. rabiei* is a necrotrophic fungus that prefers humid conditions and low temperatures (20°C). The primary infection source is airborne or water-borne conidia and ascospores (Ilarslan and Dolar, 2002; Nizam et al., 2010; Sharma and Ghosh, 2016). After penetration, fungal hyphae accumulate in cortical cells and differentiate into asexual spores called pycnidia (Pandey et al., 1987) that are responsible for secondary infection. High moisture enhances secondary infections and increases the number of spores that persists in the soil. Some areas affected by Ascochyta blight in different parts of the world are also prone to drought stress (Tadesse et al., 2017; Sinha et al., 2019), providing the rationale to study the effect of drought on fungal infection.

In the field, the combination of drought stress and fungal infection results in a complex interaction of defense pathways from both the host and the fungus, resulting in the suppression or the intensification of the infection. Drought-mediated tolerance to fungal infection may have two distinct causes. First, drought might suppress fungal growth and reproduction, thus reducing the fungal inoculum (Markel et al., 2008). Second, the defense responses elicited by drought may act as an added arsenal to protect plants from invading pathogens (Fabro et al., 2004; Chen and Dickman, 2005; Achuo et al., 2006; Ramegowda et al., 2013; Ayoubi and Soleimani, 2014). Plant responses to combined stress are themselves a combination of shared and unique molecular and physiological responses (Pandey et al., 2015). The activation of reactive oxygen species (ROS) detoxification pathways, the downregulation of the photosynthetic machinery, the upregulation of stress-responsive genes, and increased accumulation of osmoprotectants are some of the molecular responses common to drought and biotic stress imposed by pathogen infections (Achuo et al., 2006; Ramegowda et al., 2013; Ayoubi and Soleimani, 2014). Several reports have shown that proline metabolism is commonly regulated by combined drought and pathogen stresses in rice (*Oryza sativa*), chickpea, and Arabidopsis (*Arabidopsis thaliana*) (Bidzinski et al., 2016; Sinha et al., 2017; Gupta et al., 2020).

Proline is synthesized from glutamate via Δ1-pyrroline-5-carboxylate synthetase (P5CS) in the cytosol and Δ1-pyrroline-5-carboxylate reductase (P5CR) in chloroplasts. After its biosynthesis, proline is transported to mitochondria by proline transporters (PTs) and oxidized by proline dehydrogenase (ProDH) and Δ1-pyrroline-5-carboxylate dehydrogenase (P5CDH) to form glutamate, with pyrroline-5-carboxylate (P5C) as an intermediate (proline–P5C cycle; Miller et al., 2009). *ProDH* expression and ProDH activity are also induced by pathogen infection. For instance, infection with *Pseudomonas syringae* pv. tomato DC3000 *AvrRpt2* in Arabidopsis raised *ProDH1* and *ProDH2* transcript levels, leading to an oxidative burst and hypersensitive response (Cecchini et al., 2011; Monteoliva et al., 2014). Similarly, ProDH2 was upregulated by combined drought and bacterial wilt in chickpea (Sinha et al., 2017). A regulated proline metabolism is critical under combined drought and pathogen infection, as proline contributes to osmoregulation and regulates redox homeostasis under stress conditions (Kishor et al., 2005). Thus, the role of enzymes of the proline–P5C cycle, particularly ProDH, should be investigated in more detail under combined drought and fungal infection in plants.

In this study, we examined the effects of drought stress on *A. rabiei* infection in chickpea under field and greenhouse conditions. We determined the changes in proline levels and the expression of the genes involved in its biosynthetic pathway under individual and combined stresses. Our results suggest a possible role for CaProDH2 in drought-mediated resistance against *A. rabiei* infection, which we validated by miRNA-induced gene silencing (MIGS) and overexpression studies. We also characterized the effects of combined drought and pathogen stresses on *CaProDH2*-silenced and overexpression lines to reveal the role of *CaProDH2* during combined stress conditions.

## Results

### Effect of drought on *A. rabiei* infection in chickpea plants

To determine the impact of drought stress on *A. rabiei* infection in chickpea, we first studied the interaction between the two stresses in a field setting in Meerut, India. Drought and combined-stress plots had ∼50% lower soil moisture content than did control plots and plots with pathogen-infected plants (Supplemental Figure S1A). The leaves and pods of 3-month-old plants exhibited an 18% decrease in disease incidence under combined-stress treatment compared to the pathogen-only treatment (Supplemental Figure S1B). We observed a similar trend at another field location in Kanpur, India (Supplemental Figure S2G–L). When grown on potato dextrose agar (PDA) medium, infected pods showed the concentric rings characteristic of *A. rabiei* growth (Supplemental Figure S2A–E). We isolated fungal pathogens from infected leaves and confirmed their identity by PCR amplification from genomic DNA and sequencing (Supplemental Figure S2F). We submitted this isolate to the Indian Type Culture Collection, India, under ITCC No. 8839.

As a second step toward expanding our understanding of the drought–*A. rabiei* interaction, we exposed chickpea plants to individual and combined stresses under controlled conditions in a growth chamber, as described in Supplemental Figure S3A. Well-watered and drought-stressed plants maintained at 80% and 30% field capacity (FC) showed relative leaf water contents of ∼85% and ∼68%, respectively (Supplemental Figure S3B). When analyzed for the incidence of Ascochyta blight 16 days post–combined-stress treatment (DPT), plants under combined stress showed reduced blight symptoms when compared to plants only infected with the pathogen (Figure 1A). We also observed more severe blight symptoms (23.5% with a score of 5) in plants exposed to the pathogen in comparison to plants subjected to combined stresses (13.9% with a score of 5) (Figure 1B; Supplemental Figure S4). Plants exposed to combined stresses also exhibited decreased cell death (Figure 1C) and reduced fungal pathogen load (Figure 1D; Supplemental Figure S5) relative to plants only infected with the pathogen. At 21 DPT, plants infected with *A. rabiei* or exposed to combined stresses showed a 50% mortality rate; upon re-watering, plants exposed to drought, combined and pathogen stresses exhibited recovery rates of 100%, 80%, and 10%, respectively (Supplemental Figure S6). To independently confirm that any condition causing water deficit has similar inhibitory effects on *A. rabiei* infection in chickpea, we exposed seedlings growing on Murashige and Skoog (MS) medium to polyethylene glycol (PEG)-induced drought and *A. rabiei* infection individually or in combination. Again, we observed that plants exposed to combined stresses displayed reduced fungal infection, increased proline accumulation, and less electrolyte damage, as compared to plants only infected with the pathogen (Supplemental Figure S7).

**Figure 1.**
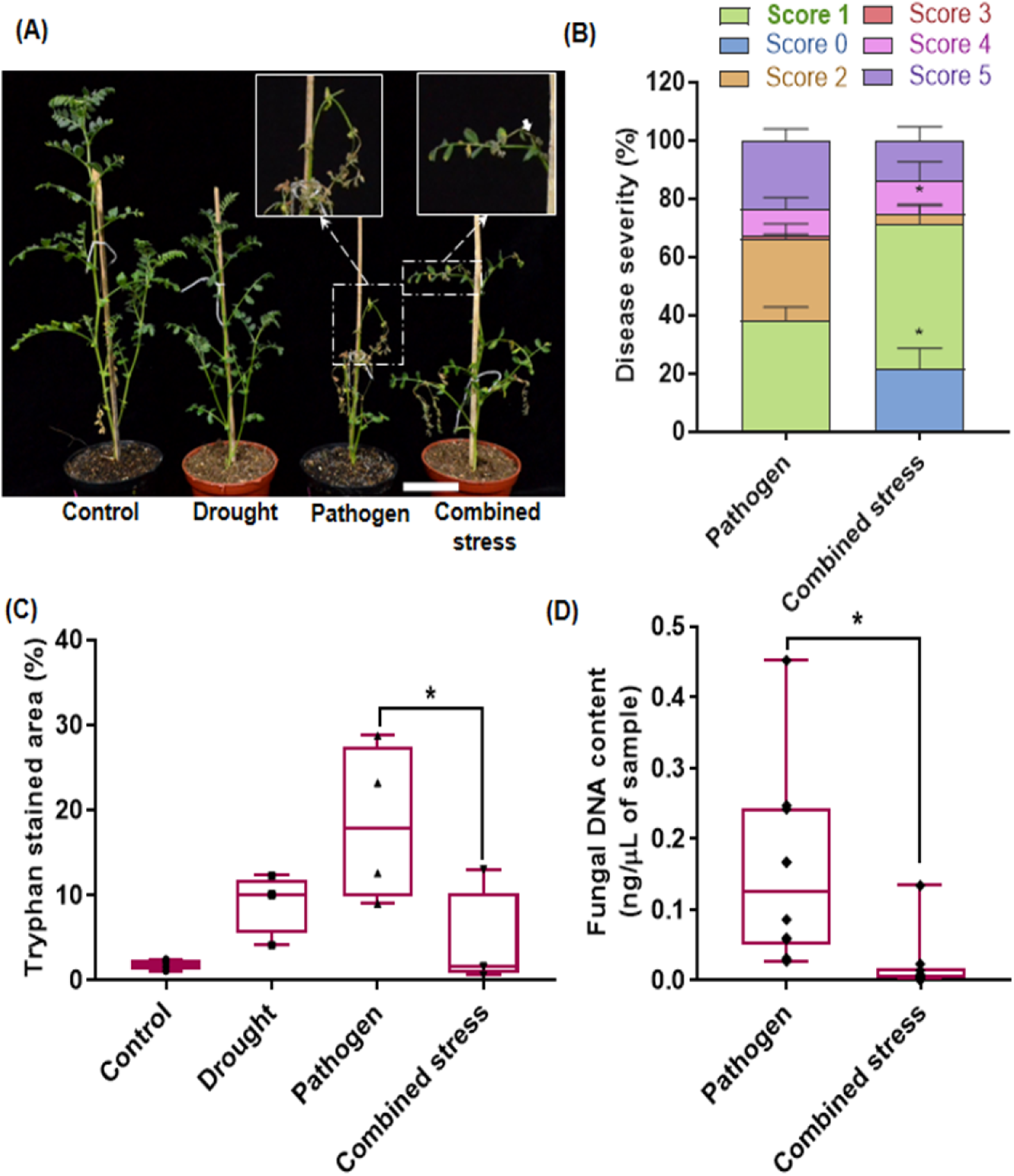
Drought imparts tolerance to *Ascochyta rabiei* infection. (A) Disease symptoms observed in chickpea plants grown in a growth room and exposed to pathogen only or combined stress in pots. Disease symptoms were documented 16 days after onset of stress. Insets, enlarged view of infected leaves, with necrotic lesions (indicated by the white arrow). (B) Assessment of disease severity in plants exposed to pathogen only or combined stresses 16 days after onset of stress, on a scale of 0 to 5. Disease scoring was carried out as described in Supplemental Figure S4. (C) Extent of cell death in infected leaves, as quantified by trypan blue–stained areas under pathogen-only or combined stresses. Quantification was performed using ImageJ software (https://imagej.nih.gov/ij/). (D) *In planta* fungal DNA content in infected leaves exposed to pathogen-only or combined stresses at 8 days after onset of stress, using *A. rabiei* specific translation elongation factor (EF) primer listed in the supplemental information. Absolute quantification values are presented here, based on the standard curve of fungal DNA shown in Supplemental Figure S5. Plant DNA of 50 ng/µL was used in qPCR. Asterisks indicate statistical significance at *p* < 0.05. The error bar indicates SEM. Experiments were repeated at least twice, with a minimum of three technical replicates. Statistical analyses were performed using one-way ANOVA (1C) and student t-test (1B, 1D), and significance is reported at *p* < 0.05.

### Effect of combined drought and *A. rabiei* infection on proline metabolism

Proline is an osmolyte involved in cell turgor maintenance; it accumulated in drought-stressed plants 7 d post–*A. rabiei* infection (Figure 2A). We also observed a concomitant increase in the levels of P5C in plants exposed to drought and combined stresses compared to controls and plants only infected with the pathogen (Figure 2A). We identified *CaProDH2* among 402 genes commonly regulated by drought and *A. rabiei* infection in a meta-analysis study of two transcriptome datasets of individual stresses (Supplemental Figures S8–S10; Supplemental File S1), prompting us to check the expression of *CaProDH2* and other proline biosynthetic genes in plants exposed to combined stresses. Drought-only and combined-stress treatments led to higher expression of genes involved in proline biosynthesis (*CaP5CS1, CaP5CS2* and *CaP5CR*), transport to mitochondria (*CaPT1*), and proline oxidation (*CaProDH2*) both at early (72 h) and late (12 DPT) stages of infection. By contrast, their expression levels decreased in plants only infected with the pathogen (Figure 2B). Further, *CaP5CDH12A1*, involved in reducing P5C to glutamate, was significantly upregulated under pathogen infection compared to combined-stress treatment (Figure 2B). These results were further confirmed by analyzing the RNAseq data available for individual and combined stresses.

**Figure 2.**
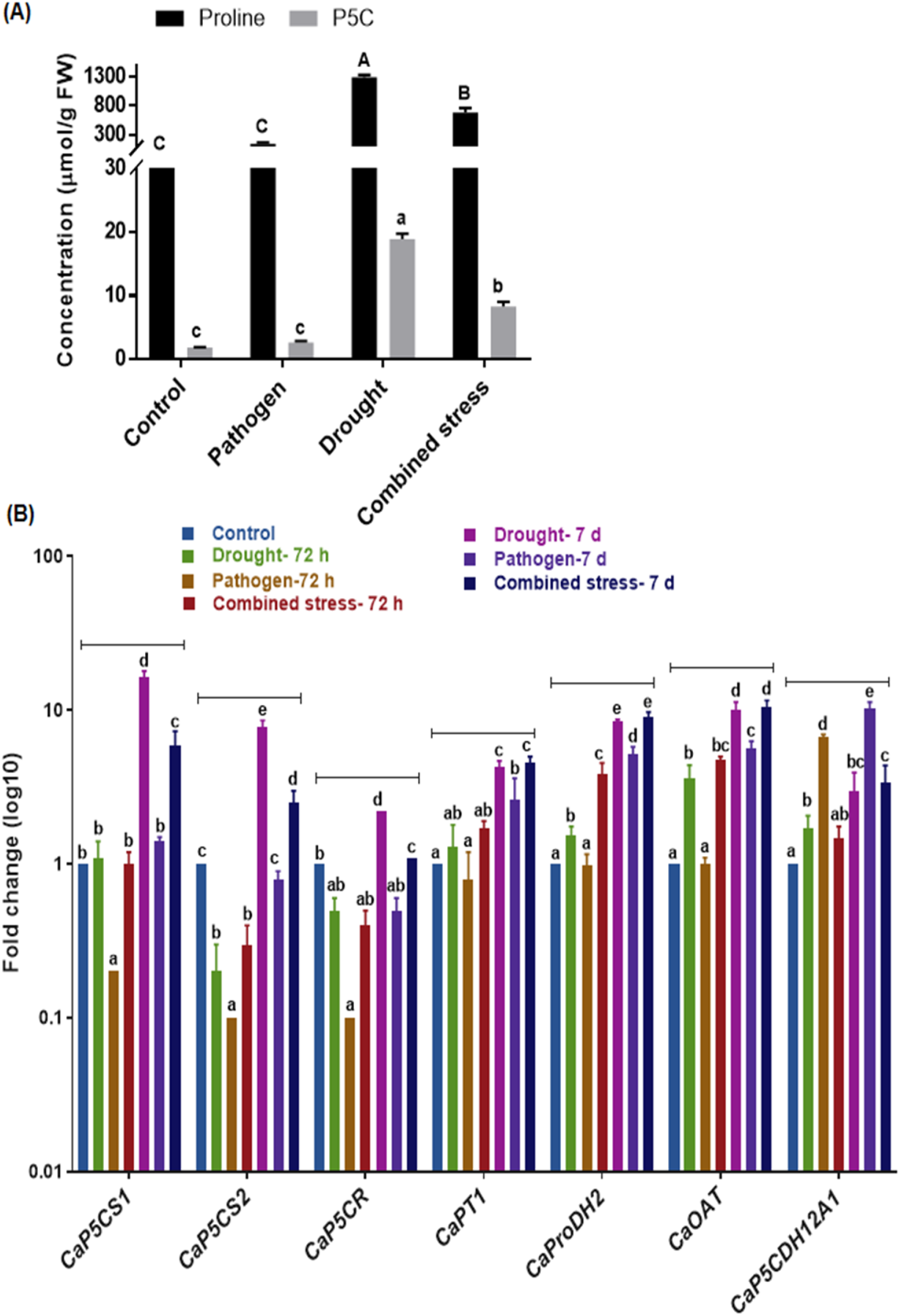
Physiological and biochemical analysis of plants subjected to combined stresses. (A) Proline and P5C content in wild-type plants subjected to different stress conditions. Chickpea plants were exposed to drought, pathogen, or combined stresses, as described in Supplemental Figure S3, and samples were collected 7 days post-infection to measure proline and P5C contents by liquid chromatography–mass spectrometry (LC-MS/MS). (B) Relative transcript levels of genes from the proline biosynthetic pathway in wild-type plants under stress, as determined by RT-qPCR. Samples were collected at 72 h and 7 d post-infection. Results are presented in log10 fold-change. The experiments were repeated three times with a minimum of three technical replicates. Different letters indicate a significant difference between treatments. Error bar indicates SEM. One-way ANOVA, followed by Duncan’s multiple range test, was performed, and significance is reported at *p* < 0.05. *P5CS, δ-1-pyrroline-5-carboxylate synthase*; *P5CR, pyrroline-5-carboxylate reductase*; *PT1, proline transporter 1*; *P5CDH, δ-1-pyrroline-5-carboxylate dehydrogenase*; *ProDH, proline dehydrogenase*; *OAT, ornithine aminotransferase*.

### Development and molecular characterization of *CaProDH2*-silenced plants

To confirm the role of *CaProDH2* in plant defense responses to combined drought and *A. rabiei* infection, we adopted the MIGS approach to knock down *CaProDH2* transcript levels (de Felippes et al., 2012) (Supplemental Figure S11). Since MIGS has not been applied in chickpea, we first tested the method by silencing a phenotypic marker gene, *PHYTOENE DESATURASE* (*CaPDS*). Accordingly, we cloned a *CaPDS* gene fragment into the MIGS2.2 vector (which also harbors an expression cassette that drives the expression of miR173 under the control of the Arabidopsis *UBIQUITIN11* promoter) and introduced it into the chickpea variety ‘Pusa 362’ (Supplemental Figures S12–S13). *CaPDS*-silenced plants showed typical photobleaching symptoms, confirming the silencing of the *CaPDS* gene and the efficacy of MIGS in chickpea (Supplemental Figures S14–S15). Results of transcriptome deep sequencing (RNA-seq) analysis of *CaPDS*-silenced plants supported the lower expression of *CaPDS* and other carotenoid biosynthesis genes without affecting genes involved in primary metabolism when miR173 is expressed (Supplemental Figure S16; Supplemental File S2).

Since MIGS appeared effective in chickpea, we generated *CaProDH2*-silenced plants using a similar strategy (Supplemental Figures S12–S13). We validated putative transgenic plants by PCR amplification of genomic DNA using *nptII*-specific primers (Figure 3). We also tested transgenic plants for antibiotic sensitivity by swabbing a kanamycin solution (150 mg/mL) on the surface of younger leaves of transformed and wild-type plants for 2 consecutive days. Only non-transgenic, wild-type plants developed bleached areas where the leaves had been exposed to the antibiotic, whereas no transgenic plant showed any bleaching (Supplemental Figure S17A). Transgenic plants that did not show signs of necrosis were selected for further analysis. These plants were healthy, with a seed-setting rate comparable to that of wild-type plants (Supplemental Figure S17D). T_2_ generation transgenics were used for assessing the stress response both at physiological and molecular level. *CaProDH2* transcript levels were analyzed in multiple lines by RT-qPCR and lines MIGS_*CaProDH2*-8, MIGS_*CaProDH2*-9, and MIGS_*CaProDH2*-12 showed significant down-regualtion of gene expression than other lines (Supplemental Figure S17B). The MIGS_*CaProDH2*-8 line was selected and used for further analysis. We confirmed the integration of the transgene in *CaProDH2*-silenced plants by Southern blot hybridization using a *nptII*-specific probe (Supplemental Figure S17C). *CaProDH2*-silenced plants expressed miR173, as evidenced by stem–loop reverse transcription–quantitative polymerase chain reaction (RT-qPCR) (Figure 3B). We detected expression of the predicted trans-acting short interfering RNAs (tasiRNAs) produced in *CaProDH2*-silenced plants via miR173-mediated silencing by stem–loop RT-qPCR using tasiRNA7- and tasiRNA12-specific primers (Figure 3C; Supplemental Figure S18). Based on *in silico* prediction of tasiRNA targets through the psRNAtarget server, *CaProDH2* was the main target of these siRNAs (Supplemental File S1). Finally, we documented miR173-mediated cleavage of *CaProDH2* by 5′ RNA Ligase-Mediated Rapid Amplification of cDNA Ends (5′ RLM-RACE), followed by sequencing of the amplified product (Figure 3D). The transgenic lines also accumulated much lower levels of endogenous *CaProDH2* transcript (Figure 3E).

**Figure 3.**
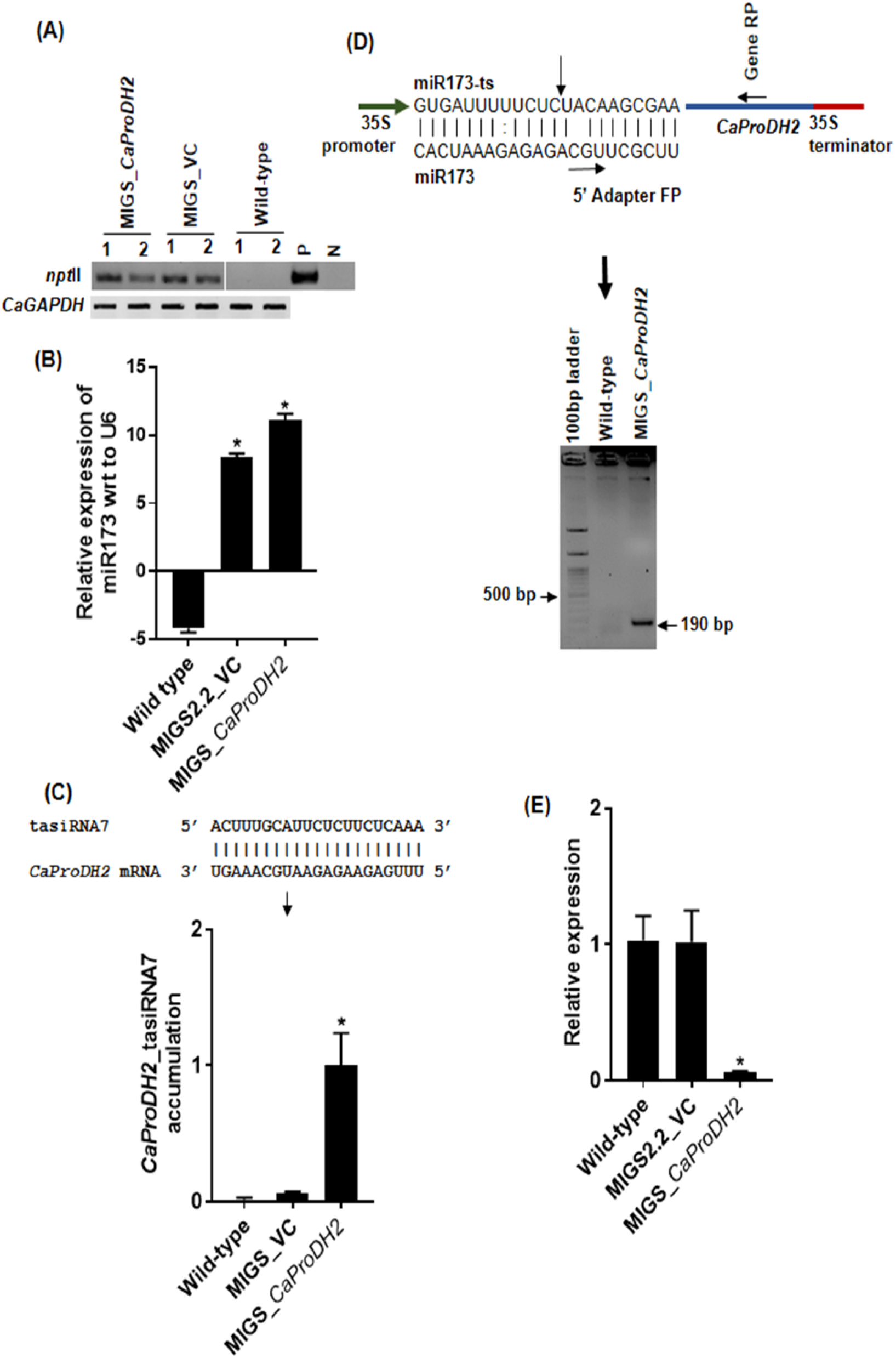
Molecular characterization of MIGS_*CaProDH2* transgenic lines. (A) Agarose gel of the *nptII* PCR amplicon in MIGS_*CaProDH2* and MIGS_VC plants. *CaGAPDH* was used as control for the PCR. Numbers 1 and 2 indicate independent transgenic events. *nptII, neomycin transferase II*; P, positive control (vector plasmid DNA); N, negative control. (B) Relative expression level of miR173, as determined by stem–loop RT-qPCR and (C) miR173-induced phased tasiRNA7 accumulation in *CaProDH2-*silenced, MIGS_VC, and wild-type plants. U6 snRNA (XR_001143939) was used as a reference. (D) Schematic representation of the miR173-binding site, 5′ adapter forward primer, and *CaProDH2* reverse primer used for 5′ RLM-RACE and confirmation of cleavage (product size 190 bp) in *CaProDH2*-silenced plants. (E) Fold-change in transcript abundance of *CaProDH2* in MIGS_*CaProDH*2, MIGS_VC, and wild-type plants. Experiments were repeated at least twice, with a minimum of three technical replicates each time. The error bar indicates SEM. Student’s *t*-test was employed, and significance is reported at *p* < 0.05.

### *A. rabiei* infection is aggravated in *CaProDH2*-silenced plants

To probe the role of *CaProDH2* in combined drought and *A. rabiei* infection, we subjected *CaProDH2*-silenced T_1_ lines to combined-stress treatment. Disease symptoms were more severe in *CaProDH2*-silenced lines than in wild-type plants under the same conditions (Figure 4A; Supplemental Figure S19A). However, both *CaProDH2*-silenced and wild-type plants were more susceptible to *A. rabiei* when only infected with the pathogen (Figure 4A). Indeed, 43% of infected leaves from *CaProDH2*-silenced lines exhibited severe blight symptoms (score 5) when well-watered, but this number decreased to 33% in combined-stress conditions. Similarly in wild-type plants, 21% or 6% of leaves showed blight symptoms when infected with the pathogen only or under combined-stress conditions, respectively (Figure 4B). *CaProDH2*-silenced plants also exhibited a more pronounced pathogen growth relative to wild-type plants during both simple infection and combined-stress conditions, as indicated by increased fungal DNA content (Figure 4C, Supplementary Figure S19B). Wild-type plants exposed to combined stress experienced the least accumulation of fungal DNA, which further supports the finding that drought suppresses Ascochyta infection in chickpea.

**Figure 4.**
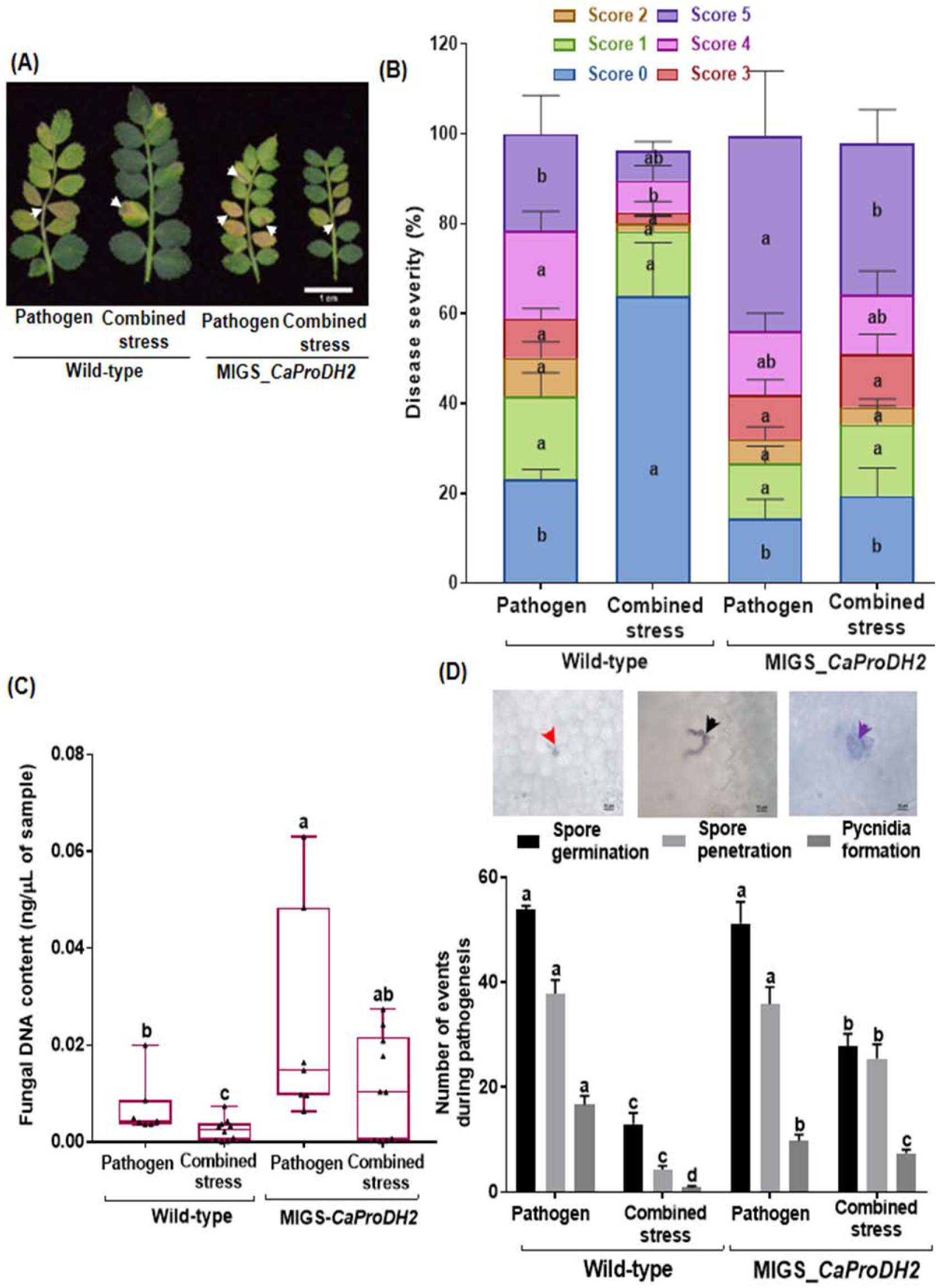
Drought-induced resistance to *Ascochyta rabiei* infection is abolished in *CaProDH2*-silenced plants. (A) Representative photographs of wild-type and *CaProDH2*-silenced plant leaves subjected to pathogen infection or combined stresses. Stress treatments were imposed as described in Supplemental Figure S3, and disease symptoms were photographed at 13 days post inoculation (DPI). White arrows indicate necrotic blight symptoms on the rachis and leaves. (B) Disease incidence and (C) fungal DNA content in plants exposed to pathogen only or to combined stresses. Disease incidence was calculated by following the method described in Supplemental Figure S4. *In planta* fungal DNA was quantified in leaf samples using a primer specific to the elongation factor gene of *A. rabiei* at 8 DPI. Absolute quantification values are presented. (D) Number of different pathogenesis events observed in wild-type and transgenic plants under pathogen or combined stresses. Infected leaves were collected and stained with trypan blue, and images at various stages of infection were captured under a 100 objective on a Nikon 80i epifluorescent microscope. Red arrow, germination of spores; black arrow, germ tube penetration; purple arrow, pycnidia formation. Different letters indicate a significant difference between the treatments and genotypes. The error bar indicates SEM. Experiments were repeated at least twice, with a minimum of three technical replicates. Two-way ANOVA, followed by Duncan’s multiple range test, was performed, and significance was reported at *p* < 0.05.

To better understand how pathogen infection and its progression are affected by drought, we carried out a detailed microscopic analysis of the infection process in wild-type and *CaProDH2*-silenced plants. We observed all three different pathogenesis stages (spore germination, germ tube elongation, and pycnidia formation; Supplemental Figures S20–S21) and counted the number of pathogenic events in both wild-type and transgenic when only infected with the pathogen or when subjected to combined stresses. Drought reduced spore germination, penetration, and pycnidia formation in both wild-type and *CaProDH2*-silenced plants (Figure 4D). The magnitude of this reduction was larger in wild-type plants exposed to combined stresses (Figure 4D). These results demonstrate a possible protective role for ProDH2 in providing defense against *A. rabiei* infection. As an additional confirmation of the role of ProDH2 in combined stress tolerance, we generated transgenic chickpea lines by transiently overexpressing *CaProDH2* and exposed them to combined drought (PEG8000, -0.9 MPa) and *A. rabiei* infection. We observed a marked increase in resistance against pathogen infection, as evidenced by reduced pathogen load in these transgenic plants (Supplemental Figure S22).

### Combined stress–induced modulation of the proline–P5C pathway in chickpea

To decipher the role of the proline–P5C cycle in defense against *A. rabiei*, we quantified the levels of proline and its oxidized form P5C in *CaProDH2*-silenced lines under control conditions, individual stress and combined stresses. *CaProDH2*-silenced lines failed to accumulate P5C, and their proline levels were 30% less than those of wild-type plants (Figure 5A, B). *CaProDH2*-silenced plants also showed a marked downregulation of proline biosynthetic genes and reduced ROS accumulation under combined stresses relative to wild-type plants (Figure 5B, 5C, Supplemental Figure S23). Furthermore, *CaProDH2*-overexpressing lines showed lesser fungal load when compared to *CaProDH2-*silenced and wild-type plants under combined stresses (Supplemental Figure S22).

**Figure 5.**
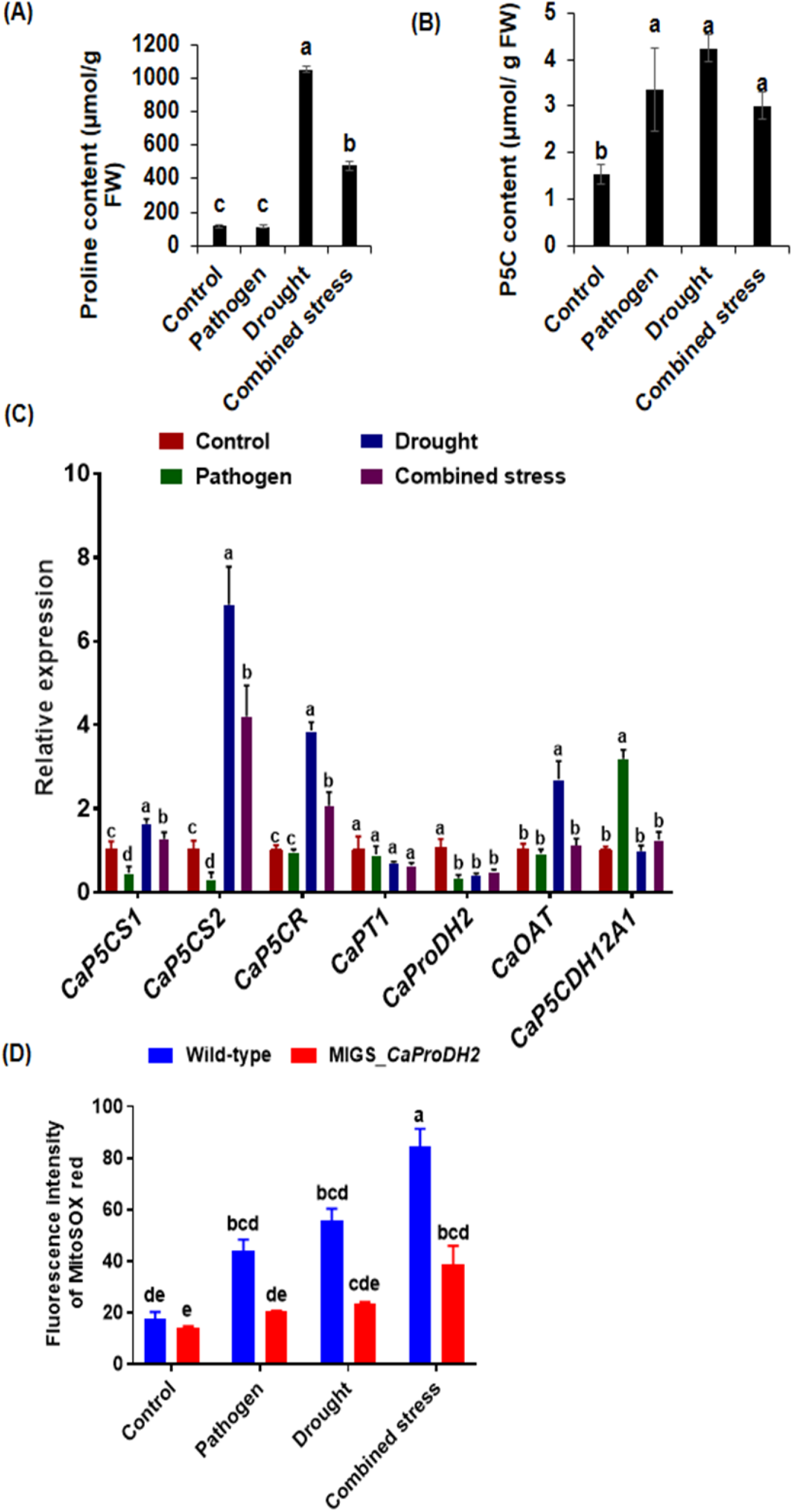
Proline, P5C, and mitochondrial ROS levels in *CaProDH2*-silenced plants. (A) Accumulation of proline and (B) P5C in wild-type and *CaProDH2*-silenced plants subjected to stress treatments. (C) Semi-quantitative expression of proline biosynthetic genes in *CaProDH2*-silenced plants. (D) Mitochondrial ROS accumulation in wild-type and *CaProDH2*-silenced plants. Different letters indicate a significant difference between the treatments and genotypes. The error bar indicates SEM. Experiments were repeated at least twice, with a minimum of three technical replicates. Two-way ANOVA, followed by Duncan’s multiple range test, was performed, and significance was reported at *p* < 0.05.

## Discussion

Our field experiments showed that drought stress decreased the severity of Ascochyta blight in chickpea. A similar drought-mediated reduction in Ascochyta blight incidence was observed in chickpea fields in Ethiopia during the drought year 2015–2016 (Tadesse et al., 2016). Pot experiments carried out in growth chambers supported observations made with field-grown plants. We established here that drought stress reduces the severity of disease symptoms and the accumulation of fungal DNA in *A. rabiei*-infected plants grown under controlled conditions. Moreover, stress-induced damage, as determined by the extent of cell death reported by Trypan blue staining, was lowest in plants experiencing combined stresses (Figure 1). Consistent with our results, low soil moisture inhibited infection by the soil-borne fungus *Sclerotium rolfsii*, which is responsible for collar rot in chickpea (Tarafdar et al., 2018). In addition, plants exposed to combined stresses showed a better recovery from drought than plants exposed to *A. rabiei* alone. Drought stress limited fungal infection and enhanced overall plant defenses, resulting in better recovery.

To investigate the mechanism behind drought-induced tolerance to *A. rabiei*, we concentrated on proline metabolism under combined drought and *A. rabiei* infection. We observed a unique modulation of proline levels and expression of its metabolism-related genes in plants exposed to combined stresses. Proline regulation is crucial in defining plant responses to both drought stress and pathogen infections (Qamar et al., 2015; Liang et al., 2013). In agreement with other reports, drought stress caused increased proline accumulation in infected chickpea plants 7 days post-inoculation. However, proline accumulation was more limited in plants subjected to combined stresses when compared to drought-stressed plants (Figure 2A). We also noticed higher expression levels of the proline catabolism gene *CaProDH2* at early and late time points following inoculation. *ProDH2* expression was also induced by *Pseudomonas syringae* pv. tomato in Arabidopsis (Cecchini et al., 2011). Furthermore, the increased levels of P5C measured in plants under combined stresses suggest that these plants metabolize proline into P5C during the entire combined stress period (Figure 2B).

We used MIGS to dissect the molecular mechanism behind drought-mediated Ascochyta blight resistance in chickpea. MIGS has been implemented in many plant species (Benstein et al., 2013; de Felippes et al., 2012; Zhou et al., 2013; Sicard et al., 2015; Zheng et al., 2018) to reduce transcript levels of various candidate genes. Here, we demonstrated the successful establishment of MIGS-mediated silencing for the first time in chickpea. As miR173 is not present in chickpea, we used the MIGS2.2 vector for its co-expression along with the tasiRNA-generating cassette, which adds the miR173 target sequence to the gene to be silenced. Transgenic plants expressing miR173 and wild-type plants were phenotypically indistinguishable. Moreover, RNA-seq analysis of wild-type plants and miR173-expressing plants revealed no marked differences between their transcriptomes, indicating that MIGS can be an effective gene silencing method in chickpea (Supplemental File S2). We then applied MIGS to generate *CaProDH2*-silenced chickpea plants, which were more susceptible to *A. rabiei* infection under both well-watered and drought-stress conditions, indicating that the loss of *CaProDH2* function compromises chickpea defense responses against the pathogen (Figure 4). Rizzi et al. (2017) reported that Arabidopsis *prodh* mutants exhibited enhanced susceptibility to the necrotrophic fungus *Botrytis cinerea. CaProDH2*-silenced lines also showed a dysregulation of proline metabolism (Figure 5). Higher ProDH activity is associated with mitochondrial superoxide production (Cecchini et al., 2011; Qamar et al., 2015). We therefore checked the levels of mitochondrial ROS in wild-type and *CaProDH2*-silenced plants. ProDH2 downregulation compromised ROS accumulation under combined stresses (Figure 5C). Likewise, the silencing of *ProDH* led to reduced ROS accumulation and compromised disease resistance in Arabidopsis (Cecchini et al., 2011; Senthil Kumar and Mysore, 2012). We showed here that the elevated P5C content in the leaves of plants experiencing combined stresses increases mitochondrial ROS production, which triggers an enhanced defense response and prevents *A. rabiei* proliferation inside chickpea. We have also validated this data with mitochondrial specific quantification of the P5C. Studies have previously reported on the role of ROS in plants defense against *Ascochyta* sp. For example, Henares et al. (2019) showed that the initial phase of *A. lentis* infection in lentils (*Lens culinaris*) results in the release of ROS. Similarly, Rea et al. (2002) showed that inhibition of H_2_O_2_ accumulation increases disease severity in lentils.

We carefully documented the various stages of fungal infection under well-watered and drought-stress conditions. Like with the *A. lentis*–lentil pathosystem, infection of *A. rabiei* in chickpea plants can be categorized into three stages: the early phase (0–4 DPI), comprising fungal adhesion, spore germination, and penetration into host cells; the mid phase (5–9 DPI), characterized by the appearance of the first symptoms due to fungal colonization of host cells; and the late stage (10 DPI onwards), showing extensive necrosis and the appearance of pycnidia on leaf tissues, indicating the onset of the following infection cycle (Pandey et al., 1987; Henares et al., 2019). We observed reduced spore germination, penetration, and pycnidia formation under combined stresses compared to plants only infected with the pathogen (Figure 4D). The *in planta* fungus transcript expression data from co-transcriptome of the chickpea infecting fungus also supported this observation. Our results agree with a previous report wherein *prodh2* mutants displayed enhanced mycelial expansion of *B. cinerea* in Arabidopsis, suggesting a role for ProDH2 in suppressing early infection events like fungal germination, penetration, and/or hyphal development (Rizzi et al., 2017). Our results showed that drought stress results in P5C accumulation and ROS induction in chickpea, which reduces fungal penetration and pycnidia formation. The reduced pycnidia formation seen in combined infected and drought-stressed plants might yield a smaller inoculum for the next infection cycle, thereby reducing the extent of leaf-to-leaf spread of the fungus (Supplemental Figure S24). Our infection experiments in transgenic plants indicated that both wild-type and *CaProDH2*-silenced plants were significantly infected by *A. rabiei*, with a comparable extent of spore germination and penetration (Figure 4). Although pycnidia formation was reduced in *CaProDH2*-silenced lines, these plants nevertheless exhibited enhanced fungal infection, as determined 13 DPI after the evaluation of the pycnidia formation stage. Reduced pycnidia formation displayed a prolonged latent phase on plants with compromised CaProDH2 function. In turn, an extended dormant period might show the extended ability of a pathogen to metabolize plant resources before forming reproductive structures (Stotz et al., 2014).

Thus, a prolonged latent phase is indicative of compromised defense responses in *CaProDH2*-silenced lines. Unlike well-watered conditions, the extent of *A. rabiei* infection was reduced under drought stress in both wild-type and *CaProDH2*-silenced plants. *CaProDH2*-silenced lines exhibited increased spore germination, penetration, and pycnidia formation compared to wild-type plants under drought stress. Loss of ProDH2 function in Arabidopsis plants accelerated the expansion of fungal mycelia, resulting in enhanced infection by *B. cinerea*, indicating a possible role for the gene in regulating the early stages of fungal infection (spore germination, penetration, and hyphal development) (Rizzi et al., 2017). Hence, reduced CaProDH2 activity, which brings a reduction in mitochondrial P5C content and ROS-mediated defense, is likely to be responsible for the higher *A. rabiei* infection in *CaProDH2*-silenced lines.

We propose that the accumulation of proline in plants exposed to both stresses induces the expression of *CaProDH2* and thus increases production of P5C. The concomitant downregulation of *P5CDH* reduces the oxidation of P5C into glutamate, consequently resulting in increased P5C levels in mitochondria. Thus, we hypothesize that after *A. rabiei* infection, the pool of proline accumulated in drought-stressed plants is oxidized to P5C in mitochondria via increased CaProDH2 levels. This conversion step and increase in mitochondrial P5C lead to ROS production, thereby enhancing defense against pathogens. The role of ProDH2 and P5C in enhancing chickpea tolerance against combined stresses is further corroborated by the observation that *CaProDH2*-silenced lines exhibit higher levels of fungal germination, maturation, and pycnidia formation than wild-type plants under combined-stress conditions (Figure 4D). Thus, we show that the accumulation of proline and the fine-tuned regulation of the proline–P5C cycle under combined stresses leads to enhanced chickpea resistance against *A. rabiei* infection under drought stress (Figure 6).

**Figure 6.**
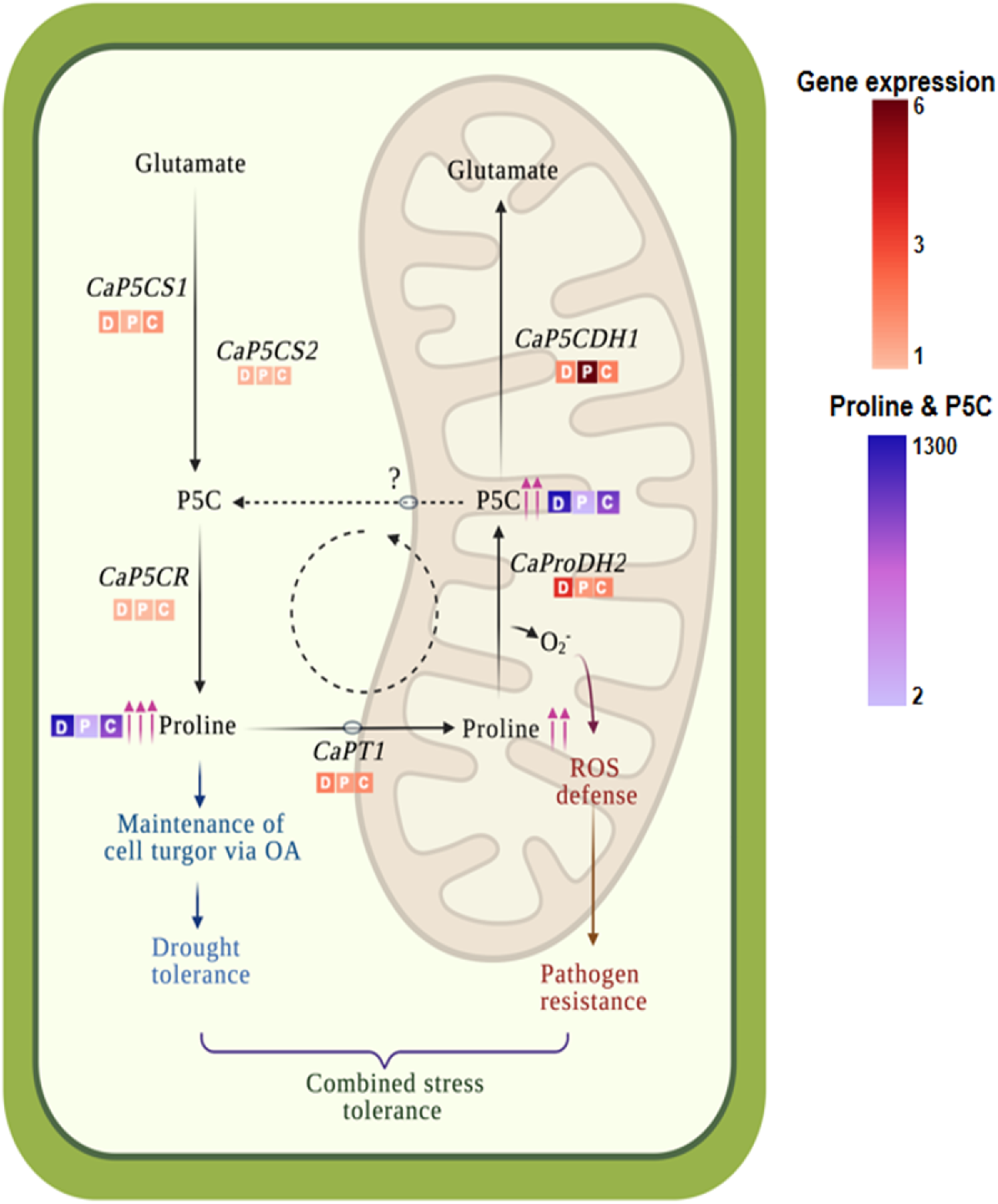
*CaProDH2*-mediated mitochondrial P5C regulation in chickpea plants subjected to combined drought and *Ascochyta rabiei* infection. Proposed model showing the regulation of the proline–P5C cycle in chickpea plants subjected to combined drought and *A. rabiei* infection. Cytosolic proline content increases in response to drought and serves as an osmolyte to maintain cell turgor, thereby conferring drought tolerance. Accumulated proline is also transported to mitochondria and converted into P5C by CaProDH2. This conversion and resulting increase in P5C level leads to the production of ROS molecules, thus inducing ROS-mediated pathogen defense responses. We observed an enhanced expression of genes involved in proline biosynthesis (*CaP5CS1, CaP5CS2, CaP5CR*), transport (*CaPT1*), and oxidization (*CaProDH2)* and higher P5C accumulation under drought and combined stress conditions. Together, increased cytosolic proline content and mitochondrial P5C with ROS enhance the resistance of chickpea plants to *A. rabiei* infection under drought stress. Blue and purple color scale, proline and P5C contents under drought (D), pathogen (P), and combined stresses (C). Red gradiant color scale, expression levels of *P5CS* (*δ-1-pyrroline-5-carboxylate synthase*); *P5CR* (*pyrroline-5-carboxylate reductase*); *PT1* (*proline transporter 1)*; *P5CDH* (*δ-1-pyrroline-5-carboxylate dehydrogenase*); and levels of P5C (1-pyrroline-5-carboxylic acid) under drought (D), pathogen (P), and combined stresses (C).

In conclusion, we report drought-induced resistance to the necrotrophic fungus *A. rabiei* in chickpea, which restricts fungal growth inside plants. Our results suggest a possible role for *CaProDH2* and the proline–P5C cycle in conferring resistance to the pathogen under drought conditions. Thus, our study highlights the dynamic role of proline metabolism under stress in chickpea. Our findings also suggest proline accumulation as an agronomically valuable trait to generate plants with higher tolerance to combined stresses. Further investigations on the role of the proline–P5C cycle in plant responses to combined stress resistance will provide the tools to breed more tolerant crops.

### Experimental procedures

#### Plant materials and growth conditions

Chickpea seeds (*Cicer arietinum* L. cv Pusa-362) were obtained from the Indian Agricultural Research Institute, New Delhi, and raised in pots (3 inches × 3 inches) containing 30 g air-dried peat and vermiculite (3:1 mixture [v/v]) in a plant growth chamber (PGR15; Conviron, Winnipeg, Canada) with a 12-h light/12-h dark cycle, 200 μE m^−2^ s^−1^ photon flux intensity, 22°C temperature, and 75% relative humidity (RH).

#### Cloning and construct development

For the validation of MIGS in chickpea, a phenotypic marker gene, *PHYTOENE DESATURASE* (*CaPDS*; XP_012571841.1), was cloned into MIGS vector MIGS2.2 (FF573) obtained from Addgene (plasmid #35248; http://n2t.net/addgene:35248; RRID: Addgene_35248). Before cloning, the gene sequence was scanned through pssRNAit (https://plantgrn.noble.org/pssRNAit/), and the regions yielding the smallest number or potential off-targets were identified. The corresponding gene segment (1,045–1,314 bp cDNA for *CaPDS*; Supplemental Table S4) was amplified from the chickpea genome by PCR and cloned into the MIGS2.2 vector using a two-step Gateway cloning method. The first step involved cloning the *CaPDS* segment in between the attL1–L2 region of the entry vector (EV1, obtained from Dr MK Reddy, ICGEB, New Delhi, India) using conventional restriction digests and was followed by recombining the gene fragment into the MIGS2.2 vector by LR clonase (Thermo Fisher Scientific, Waltham, MA, USA) (Supplemental Figure S10). Cloning was confirmed by PCR amplification and sequencing with gene-specific primers (Supplemental Table S5). The resulting construct was transformed into Agrobacterium (*Agrobacterium tumefaciens*) strain GV3101 using the freeze–thaw method. As MIGS2.2 has a pGreen backbone, pSOUP was co-transformed along with the MIGS vector. A 259-bp *C. arietinum PROLINE DEHYDROGENASE2* (*CaProDH2*; 386–644 bp for *CaProDH2* GenBank Acc. No. XM_004491715; Supplemental Table S4) gene fragment was also cloned into the MIGS vector using the same steps as above. The primers used for cloning are listed in Supplemental Table S5.

### Preparation of Agrobacterium cultures for plant transformation

#### Plant transformation

Chickpea transformation was performed according to Sarmah et al. (2004) and Khandal et al. (2020), with a few modifications. Mature disease-free chickpea seeds (*C. arietinum* cv. Pusa-362) were surface-sterilized with ethanol (70% for 1 min) followed by mercuric chloride (0.1% for 10 min) and rinsed three to four times in sterile distilled water before soaking in sterile distilled water overnight. The primary Agrobacterium inoculum was prepared by inoculating a single colony harboring pMIGS2.2_CaPDS, pMIGS2.2_CaProDH2, or empty vector (pMIGS2.2 without the CcdB region) in 5 mL Luria Bertani (LB; HiMedia Laboratories, Mumbai, India) medium containing 50 mg/L each spectinomycin and rifampicin. Flasks were incubated at 28°C with shaking at 180 rpm overnight. Secondary cultures were started by inoculating 50 mL LB medium with 500 µL primary inoculum. Agrobacterium cells (OD600 = 0.6–0.8) were harvested by centrifugation at 2,960 g for 10 min in a hybrid refrigerated centrifuge (CAX-371; Tomy, Tokyo, Japan). Cell pellets were resuspended in 50 mL Agrobacterium induction medium, which was prepared by adding 2.5 mL 1 M MES-KOH buffer pH 5.7 (HiMedia Laboratories), 100 µL 1 M MgCl_2_ (Fisher Scientific, Newington, USA), 250 mg glucose (Amresco, Solon, OH, USA) and 100 µM acetosyringone (HiMedia Laboratories) and incubated at 25°C with shaking at 80 rpm for 3 h. The seed coat was removed and each chickpea seed was then dissected into two halves, each with one cotyledon and half of the embryonic stem, in the presence of a bacterium inoculum. The half embryos were incubated with the bacterial culture at room temperature for another 15 min, blot-dried, and transferred to full-strength MS medium, pH 5.8 (Duchefa Biochemie, Haarlem, the Netherlands) (Murashige and Skoog, 1962) with 3% sucrose (Fisher Scientific) and 0.8% agar. After 2 days of co-cultivation, explants with green shoots were transferred to the kanamycin selection medium. Sub-culturing was performed four times with a gradual increase in antibiotic concentrations (50, 100, 150, and 200 mg/L). Plantlets were subjected to hardening for soil establishment and grown at 20°C in a long-day photoperiod with 150 μmol m^-2^ s^-1^ light intensity and 55% ± 2% of RH. Fully grown plants were used for experiments. The presence of the transgene was confirmed by PCR on genomic DNA using *npt*II-specific primers. Genomic DNA was isolated from the leaves of vector control plants and transformants by using DNAzol (Invitrogen, California, USA) solution and following the manufacturer’s protocol.

#### RT-qPCR and stem–loop RT-PCR

Total RNA from control and stressed samples of wild-type and *CaProDH2*-silenced plants was isolated using TRIzol reagent. First-strand cDNAs were synthesized from 2 µg of total RNA and analyzed for gene expression by RT-qPCR following the protocol described by Gupta et al., 2020. Details of the methodology followed for RT-qPCR and stem–loop RT-qPCR are given in Supplemental Methods S1. The primers for stem–loop RT-qPCR were designed according to Chen et al. (2005a) (Supplemental Table S5).

### Stress treatment

#### Preparation of fungal inoculum and plant infection

The details of *A. rabiei* inoculation are described in an earlier publication (Verma et al., 2017). The mini-dome technique described by Chen et al. (2005b) was used for *A. rabiei* infection of chickpea plants. Single-spore isolates of *A. rabiei* (ITCC-4638) were grown on PDA medium (HiMedia Laboratories). Spores were released from petri plates containing PDA-grown fungal cultures by adding 2 mL sterile water and incubating for 15 min with frequent scraping using a sterile loop. After the suspension was filtered through muslin cloth, the spore titer was determined on a hemocytometer. The spore suspension was diluted to 1 × 10^6^ spores mL^−1^ in sterile water. Control and drought-stressed plants (21 days old) were sprayed with the spore suspension containing 0.05% Tween-20 (HiMedia Laboratories) and 0.1% of sucrose to reduce run-off. Plants were covered with inverted translucent plastic cups to form a mini-dome for 5–6 days to maintain high humidity. Plants were then placed in a growth chamber set to 22°C ± 2°C and 70% ± 5% RH with a photoperiod of 14-h light/10-h dark. Symptoms were recorded 7–10 days post inoculation (DPI, Supplemental Figure S3).

#### Drought stress

Pre-weighed 15-day-old chickpea seedlings were subjected to drought stress by the gravimetric approach (Ramegowda et al., 2013). Twice a day, pots were weighed until soil moisture content reached 30% FC (Ψw - 1.0 MPa), after which point pots were maintained at 30% FC until the end of the experiment. Control plants were maintained at 80% FC (Ψw - 0.8 MPa) by replenishing the amount of water lost twice a day throughout the course of the experiment. The vapor pressure deficit in the growth chamber was 0.793 kPa. The soil moisture content (i.e., FC) was calculated using the following formula, where WW is wet weight and DW is dry weight:

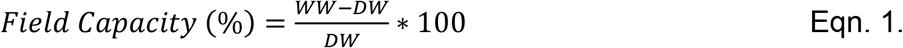

#### Combined stress treatment

Combined stresses were applied by exposing drought-stressed plants maintained at 30% FC to *A. rabiei* infection. The mouth of the pots was covered with cling wrap to avoid any water dripping onto the soil as a result of spraying. A spore suspension of 1 × 10^6^ spores mL^−1^ was sprayed on drought-stressed chickpea plants until run-off. Mock-infected control and drought-stressed plants were sprayed with sterile water and covered with transparent plastic sheets to create high humidity. Pots were weighed after spraying to make sure that FC was unchanged. The timeline of combined stress treatment is depicted in Supplemental Figure S3.

### Determination of proline and P5C content

The leaves of chickpea plants subjected to control, drought, *A. rabiei*, and combined stresses were used for proline and P5C quantification. Proline content was estimated using a published protocol by Bates (1973). Tissue samples 12 days into stress treatment were ground in 2 mL 3% aqueous sulfosalicylic acid (Fisher Scientific) solution. The homogenate was filtered using whatman filter paper and mixed with 2 mL acid ninhydrin (1.25 g ninhydrin [Fisher Scientific] + 30 mL glacial acetic acid [Merck, New Jersey, USA] + 20 mL 6 M phosphoric acid [Fisher Scientific]) and 2 mL of glacial acetic acid in sterile test tubes. Tubes were heated to 100°C for 1 h and then transferred to an ice water bath to terminate the reaction. To each tube, 4 mL toluene (Fisher Scientific) was added and mixed vigorously. The upper toluene supernatant fraction was taken, and the absorbance was recorded at 520 nm. Using a proline standard curve, the concentration of proline in samples was determined and expressed on a fresh weight basis. For the standard curve, 10, 20, 40, 60, 80, and 100 µg proline was prepared, and absorbance was recorded at 520 nm (the standard graph is shown in Supplemental Figure S5). Also using liquid chromatography–mass spectrometry (LC-MS/MS) proline and P5C content were measured from stressed samples. Details of the methodology followed for GC-MS analysis are given in Supplemental Methods S1.

### Fungal DNA estimation

Fungal DNA was quantified from plant samples according to Bayrakatar et al. (2016). Fungal genomic DNA was isolated from an *A. rabiei* culture grown at 20°C on PDA medium. For DNA extraction, cultivated mycelia (100 mg) or plant tissue samples infected with *A. rabiei* were used, and DNA was isolated using the DNAzol reagent. Using pure fungal DNA (100–0.001 ng), a standard curve was obtained by qPCR, as described above, using *HEF* forward and reverse primers (450 nM). The standard curve was generated by plotting Ct values against the known fungal DNA concentration. Similarly, using the genomic DNA isolated from plant samples (diluted to 10 and 50 ng L^−1^), qPCR was performed and the concentration of fungal DNA determined from the Ct values and the standard curve.

### Microscopy observations

Leaf and stem samples were collected and observed for blight lesions, cell death, fungal spores, and fungal infection under either 40 × or 100× magnification using a Nikon Eclipse 80i epifluorescence microscope (Nikon Corporation, Tokyo, Japan) equipped with a Nikon digital camera (Nikon Digital Sight DSFi3, New York, USA) or using a Nikon AZ100 stereo fluorescence microscope with a 0.5× objective lens equipped with a Nikon digital camera (Nikon Digital Sight DS-Ri1).

### mRNA cleavage assay

The miRNA173-mediated cleavage of *CaProDH2* was confirmed by 5′ RLM-RACE. Total RNA from leaf tissue (100 mg, 4-week-old plants) was extracted by using the TRIzol reagent. mRNA isolation, purification, and 5′ adapter ligation were performed according to Gupta et al. (2020). The resulting products were amplified using a 5′ adapter forward and 3′ reverse gene-specific primer, followed by confirmation of amplified fragments by sequencing and multiple sequence alignment.

The methodology followed for the other experiments is given in Supplemental Methods S1.

### Mitochondrial ROS estimation

Mitochondrial superoxide radicals were quantified according to Cvetkovska and Vanlerberghe (2013). Leaves from wild-type and *CaProDH2*-silenced plants subjected to drought, *A. rabiei* infection, and combined stresses were treated with 1% dimethyl sulfoxide (DMSO) for 5 min, followed by floating in 3 mM MitoSOX Red (Catalog# M36008, Fisher Scientific) and 0.35 mM MitoTracker Red CMXRos (Catalog# M7512, Fisher Scientific) solution in the dark for 30 and 20 min respectively at 37 °C. Samples were then removed from the solution, washed with sterile water and mounted onto slides and examined on a Laser Scanning Microscope (AOBS TCS-SP5, LEICA GERMANY) with appropriate excitation/detection settings (MitoSOX Red, 488/585–615 nm; MitoTracker Red, 543/585–615 nm).

## Statistical analysis

Data presented in this manuscript were analyzed using GraphPad Prism 7 (https://graphpad.com/scientific-software/prism/) and MSTAT-C (https://msu.edu/~freed/mstatc.htm) software. Significant differences between genotypes and across the treatments were tested by ANOVA, followed by Duncan’s multiple range test (DMRT) (*p <* 0.05). Raw data of all figures and tables presented in this manuscript are given in Supplemental File S3.

## Accession numbers

PDS (XM_012716387), ProDH2 (XM_004491715).

## Acknowledgments

Projects in the MS-K lab are supported by the National Institute of Plant Genome Research core funding. PP, MP, and VI acknowledge SERB (SERB/LS-359/2014), CSIR (No.13 (9064-A)/2019-Pool), and DBT-JRF (DBT/2015/NIPGR/430), respectively. We thank Dr Sandhya Verma and Ms Ankita Shree for their help in carrying out the fungal infections. We acknowledge Dr Debasis Chattopadhyay and Dr Santosh Kumar Gupta for providing the chickpea transformation protocol and transgenic facility for plant genetic transformation support. We also thank Ms Anju Udhay, Mr Pandiyan Muthuramalingam, Mr Masthan Basha and Mr Sombir Rao for their technical help. We also thank Mr Sundar, Mr Rahim, Mr Prem Negi and Mr Ashok Kumar for technical help at the field and central instrumentation facility. We acknowledge DBT-eLibrary Consortium (DeLCON) and NIPGR library for providing access to e-resources and NIPGR Plant Growth Facility for plant growth support.

## Legends for supporting information

**Supplemental File S1**. List of differentially expressed genes (DEGs) common to drought stress and *Ascochyta* infection, as analyzed by SAM and INMEX, and targets of predicted tasiRNAs resulting from the expression of the MIGS_*CaProDH2* construct, as analyzed by psRNAtarget.

**Supplemental File S2**. List of DEGs in MIGS_*CaPDS* transgenic plants relative to wild-type plants.

**Supplemental File S3**. Raw data of all figures presented in this study.

**Supplemental Figure S1**. Incidence of Ascochyta blight under well-watered and drought-stress conditions in the field.

**Supplemental Figure S2**. Ascochyta blight (AB) disease symptoms in field conditions, isolation of *A. rabiei* and confirmation of AB disease in chickpea fields.

**Supplemental Figure S3**. Details of protocol followed to impose combined stress and analysis of stress response.

**Supplemental Figure S4**. Assessment of disease severity in plants subjected to pathogen only and combined-stress treatments.

**Supplemental Figure S5**. Standard curve used to determine the extent of fungal infection in stressed samples using pure fungal genomic DNA.

**Supplemental Figure S6**. Response of chickpea plants to drought recovery.

**Supplemental Figure S7**. Establishment of MS medium-based combined stress imposition protocol.

**Supplemental Figure S8**. Flow chart depicting the steps involved in the meta-analysis of transcriptomic data under individual drought or *A. rabiei* infection stress.

**Supplemental Figure S9**. Meta-analysis of transcript levels under individual drought or *A. rabiei* infection stress.

**Supplemental Figure S10**. Transcriptome profiling of genes selected from meta-analysis.

**Supplemental Figure S11**. Schematic representation of miR173-mediated generation of tasiRNAs in plants.

**Supplemental Figure S12**. Schematic representation of the vectors and constructs used in this study.

**Supplemental Figure S13**. Methodology followed for the development of chickpea transformation.

**Supplemental Figure S14**. Analysis of MIGS_*CaPDS* transgenic plants.

**Supplemental Figure S15**. Schematic representation of miR173-mediated cleavage and generation of predicted tasiRNAs.

**Supplemental Figure S16**. Effect of *CaPDS* silencing on the carotenoid biosynthetic pathway.

**Supplemental Figure S17**. Assessment of kanamycin sensitivity, *CaProDH2* gene expression, and southern analysis of MIGS_*CaProDH2* transgenic plants.

**Supplemental Figure S18**. Expression of tasiRNAs in transgenic and vector control plants.

**Supplemental Figure S19**. Assessment of combined stress responses in MIGS_*CaProDH2* transgenic plants.

**Supplemental Figure S20**. Stages of *A. rabiei* infection in chickpea.

**Supplemental Figure S21**. Schematic representation of the different stages of *A. rabiei* infection in chickpea.

**Supplemental Figure S22**. Analysis of transient overexpressed *CaProDH2* plants.

**Supplemental Figure S23**. Superoxide production and colocalization with mitochondria in *CaProDH2*-silenced and wild-type plants.

**Supplemental Figure S24**. Model of the influence of drought stress on *A. rabiei* soil inoculum and plant infection under field conditions.

**Supplemental Table S1**. List of studies on the effect of drought or *A. rabiei* infection on the transcriptome of chickpea plants.

**Supplemental Table S2**. List of DEGs commonly shared by drought and *A. rabiei* infection, as determined through meta-analysis of microarray data sets of individual responses to drought or *A. rabiei* infection.

**Supplemental Table S3**. List of genes involved in photosynthesis, light signaling, and cell cycle, with their expression estimates in the MIGS (vector control) line, as compared to wild-type.

**Supplemental Table S4**. Details of marker and target genes selected for cloning into the MIGS2.2 vector.

**Supplemental Table S5**. List of primers used in this study.

## Parsed Citations

Achuo, E.A., Prinsen, E. and Höfte, M. (2006) Influence of drought, salt stress and abscisic acid on the resistance of tomato to Botrytis cinerea and Oidium neolycopersici. Plant Pathol. 55, 178–186

Ayoubi, N. and Soleimani M.J. (2014) Possible effects of pathogen inoculation and salicylic acid pre-treatment on the biochemical changes and proline accumulation in green bean. Arch. Phytopathol. Plant Prot. 48, 212–222

Bates, L.S., Waldren, R.P. and Teare, I.D. (1973) Rapid determination of free proline for water-stress studies. Plant Soil 39, 205–207

Bayraktar, H., Ozer, G., Aydoğan, A. and Palacioglu, G. (2016) Determination of Ascochyta blight disease in chickpea using real-time PCR. J Plant Dis Prot. 123, 109–117

Benstein, R. M., Ludewig, K., Wulfert, S., Wittek, S., Gigolashvili, T., Frerigmann, H., Gierth, M., Flügge, U. I., and Krueger, S. (2013) Arabidopsis phosphoglycerate dehydrogenase1 of the phosphoserine pathway is essential for development and required for ammonium assimilation and tryptophan biosynthesis. Plant Cell 25(12), 5011–5029

Bhatti, M.A. and Kraft, J.M. (1992) Effects of inoculum density and temperature on root rot and wilt of chickpea. Plant Dis. 76, 50–54

Bidzinski, P., Ballini, E., Ducasse, A., Michel, C., Zuluaga, P., Genga, A., Chiozzotto, R. and Morel, J. -B. (2016) Transcriptional Basis of Drought-Induced Susceptibility to the Rice Blast Fungus Magnaporthe oryzae. Front. Plant Sci. 7, 1558

Cecchini N. M., Monteoliva M. I. and Alvarez M. E. (2011) Proline dehydrogenase contributes to pathogen defense in Arabidopsis. Plant Physiol. 155, 1947–1959

Chen, C. and Dickman, M. B. (2005). Proline suppresses apoptosis in the fungal pathogen Colletotrichum trifolii. Proc. Natl. Acad. Sci. U.S.A. 102, 3459–3464.

Chen, C., Ridzon, D. A., Broomer, A. J., Zhou, Z., Lee, D. H., Nguyen, J. T., Barbisin, M., Xu, N. L., Mahuvakar, V. R., Andersen, M. R., Lao, K. Q., Livak, K. J., & Guegler, K. J. (2005a) Real-time quantification of microRNAs by stem-loop RT-PCR. Nucleic Acids Res. 33(20), e179.

Chen, W., Mcphee, K.E. and Muehlbauer, F.J. (2005b). Use of a mini-dome bioassay and grafting to study resistance of chickpea to Ascochyta blight. J. Phytopathol. 153, 579–587.

Coram, T.E. and Pang, E.C. (2006) Expression profiling of chickpea genes differentially regulated during a resistance response to Ascochyta rabiei. Plant Biotechnol. J. 4, 647– 666.

Conesa, A., and Götz, S. (2008). Blast2GO: Acomprehensive suite for functional analysis in plant genomics. Int. J. Plant Genomics, 619832.

Cvetkovska M, Vanlerberghe GC. (2012) Alternative oxidase modulates leaf mitochondrial concentrations of superoxide and nitric oxide. New Phytologist 195(1),32–39.

de Felippes, F. F., Wang, J. and Weigel, D. (2012) MIGS: miRNA-induced gene silencing. Plant J. 70, 541– 547.

Fabro, G., Kovacs, I., Pavet, V., Szabados, L. and Alvarez, M. E. (2004). Proline accumulation and AtP5CS2 gene activation are induced by plant-pathogen incompatible interactions in Arabidopsis. Mol. Plant Microbe Interact. 17, 343–350.

Gupta, A., Patil, M., Qamar, A., and Senthil-Kumar M. (2020) ath-miR164c influences plant responses to the combined stress of drought and bacterial infection by regulating proline metabolism. Environ. Exp. Bot. 172, 103998

Han, Y., Zhang, B., Qin, X., Li, M. and Guo, Y. (2015) Investigation of a miRNA-Induced Gene Silencing Technique in Petunia Reveals Alterations in miR173 Precursor Processing and the Accumulation of Secondary siRNAs from Endogenous Genes. PLoS ONE 10(12), e0144909

Henares, B. M., Debler, J. W., Farfan-Caceres, L. M., Grime, C. R., and Lee, R. C. (2019) Agrobacterium tumefaciens-mediated transformation and expression of GFP in Ascochyta lentis to characterize ascochyta blight disease progression in lentil. PloS One 14(10), e0223419

Ilarslan, H. and Dolar, F.S. (2002) Histological and Ultrastructural Changes in Leaves and Stems of Resistant and Susceptible Chickpea Cultivars to Ascochyta rabiei. J. Phytopathol. 150, 340–348

Jaiswal, P., Cheruku, J.R., Kumar, K., Yadav, S., Singh, A. Kumari, P., Dube, S.C., Upadhyaya, K.C. and Verma, P.K. (2012) Differential transcript accumulation in chickpea during early phases of compatible interaction with a necrotrophic fungus Ascochyta rabiei. Mol. Biol. Rep. 39, 4635–4646

Khandal, H., Gupta, S.K., Dwivedi, V., Mandal, D., Sharma, N.K., Vishwakarma, N.K., Pal, L., Choudhary, M., Francis, A., Malakar, P., Singh, N.P., Sharma, K., Sinharoy, S., Singh, N.P., Sharma, R. and Chattopadhyay, D. (2020) Root-specific expression of chickpea cytokinin oxidase/dehydrogenase 6 leads to enhanced root growth, drought tolerance and yield without compromising nodulation. Plant Biotechnol. J. DOI: https://doi.org/10.1111/pbi.13378.

Kishor, P.K., Sangam, S., Amrutha, R., Laxmi, P.S., Naidu, K., Rao, K., Rao, S., Reddy, K., Theriappan, P. and Sreenivasulu, N. (2005) Regulation of proline biosynthesis, degradation, uptake and transport in higher plants: its implications in plant growth and abiotic stress tolerance. Curr. Sci. 88, 424–438

Liang, X., Zhang, L., Natarajan, S. K., and Becker, D. F. (2013) Proline mechanisms of stress survival. Antioxid. Redox Signal. 19(9), 998– 1011

Mantri, N. L., Ford, R., Coram, T. E., and Pang, E. C. (2007) Transcriptional profiling of chickpea genes differentially regulated in response to high-salinity, cold and drought. BMC Genomics 8, 303

Markell S, Khan M, Secor G, Gulya T, Lamey A (2008) Row crop diseases in drought years NSDU-PP1371. http://www.ag.ndsu.edu/publications/landing-pages/crops/row-crop-diseasesin-drought-years-pp-1371.

Miller, G., Honig, A., Stein, H., Suzuki, N., Mitler, R. and Zilberstein, A. (2009) Unraveling delta1-pyrroline-5-carboxylate-proline cycle in plants by uncoupled expression of proline oxidation enzymes. J. Biol. Chem. 289, 26482–26492

Monteoliva, M. I., Rizzi, Y. S., Cecchini, N. M., Hajirezaei, M. R. and Alvarez M. E. (2014) Context of action of proline dehydrogenase (ProDH) in the hypersensitive response of Arabidopsis. BMC Plant Biol. 14, 21

Nizam, S., Singh, K. and Verma, P.K. (2010) Expression of the fluorescent proteins DsRed and EGFP to visualize early events of colonization of the chickpea blight fungus Ascochyta rabiei. Curr Genet. 56, 391–399

Pande, S., Siddique, K.H.M., Kishore, G.K. et al., 2005. Ascochyta blight of chickpea (Cicer arietinum L.): a review of biology, pathogenicity, and disease management. Aust. J. Agric. Res. 56, 317–32

Pandey, B.K., Singh, U.S., and Chaube, H.S. (1987) Mode of infection of Ascochyta blight as caused by Ascochyta rabiei. J. Phytopathol. (Berlin) 119, 88–93

Pandey, P., Ramegowda, V., and Senthil-Kumar, M. (2015) Shared and unique responses of plants to multiple individual stresses and stress combinations: physiological and molecular mechanisms. Front. Plant Sci. 6,723.

Pandey, P., Irulappan, V., Bagavathiannan, M.V. and Senthil-Kumar, M. (2017) Impact of combined abiotic and biotic stresses on plant growth and avenues for crop improvement by exploiting physio-morphological traits. Front. Plant Sci. 8, 537.

Qamar, A., Mysore, K. S., and Senthil-Kumar, M. (2015). Role of proline and pyrroline-5-carboxylate metabolism in plant defense against invading pathogens. Front. Plant Sci. 6, 503.

Ramegowda, V. and Senthil-Kumar, M. (2015) The interactive effects of simultaneous biotic and abiotic stresses on plants: mechanistic understanding from drought and pathogen combination. J. Plant Physiol. 176, 47–54.

Ramegowda, V., Senthil-Kumar, M., Ishiga, Y., Kaundal, A., Udayakumar, M. and Mysore, K. S. (2013). Drought stress acclimation imparts tolerance to Sclerotinia sclerotiorum and Pseudomonas syringae in Nicotiana benthamiana. Int. J. Mol. Sci. 14, 9497–9513.

Rea, G., Metoui, O., Infantino, A., Federico, R., Angelini, R. (2002). Copper amine oxidase expression in defense responses to wounding and Ascochyta rabiei invasion. Plant Physiol. 128(3), 865–875.

Rizzi, Y. S., Cecchini, N. M., Fabro, G., and Alvarez, M. E. (2017). Differential control and function of Arabidopsis ProDH1 and ProDH2 genes on infection with biotrophic and necrotrophic pathogens. Mol Plant Pathol. 18(8), 1164–1174.

Sarmah, B. K., Moore, A., Tate, W., Molvig, L., Morton, R. L., and Rees, D. P. (2004) Transgenic chickpea seeds expressing high levels of a bean α-amylase inhibitor. Mol. Breed. 14, 73–82.

Sharma, M. and Ghosh, R. (2016) An update on genetic resistance of chickpea to Ascochyta blight. Agronomy 6, 18

Sharma, M., and Pande, S. (2013) Unravelling effects of temperature and soil moisture stress response on development of dry root rot [Rhizoctonia bataticola (Taub.)] Butler in Chickpea. Am. J. Plant Sci. 4, 584–589

Sicard, A., Kappel, C., Josephs, E.B., Lee, Y.W., Marona, C., Stinchcombe J.R., Wright, S.I. and Lenhard M. (2015). Divergent sorting of a balanced ancestral polymorphism underlies the establishment of gene-flow barriers in Capsella. Nat. Commun. 6, 7960

Sinha, R., Gupta, A., and Senthil-Kumar, M. (2017) Concurrent drought stress and vascular pathogen infection induce common and distinct transcriptomic responses in chickpea. Front. Plant Sci. 8, 333.

Sinha, R., Irulappan, V., Mohan-Raju, B., Suganthi, A., and Senthil-Kumar, M. (2019) Impact of drought stress on simultaneously occurring pathogen infection in field-grown chickpea. Scientific Rep. 9(1), 5577.

Stotz, H.U., Mitrousia, G.K., de Wit, P.J.G.M. and Fitt, B.D.L.(2014) Effector-triggered defence against apoplastic fungal pathogens. Trends Plant Sci. 19, 491–500

Tadesse, M., Turoop, L., and Ojiewo, C.O. (2017) Survey of Chickpea (Cicer arietinum L) Ascochyta Blight (Ascochyta rabiei Pass.) Disease Status in Production Regions of Ethiopia. Plant 5(1), 23.

Tarafdar, A., Rani, T.S., Chandran, U.S.S., Ghosh, R., Chobe, D.R. and Sharma, M. (2018) Exploring combined effect of abiotic (soil moisture) and biotic (Sclerotium rolfsii Sacc.) stress on collar rot development in chickpea. Front. Plant Sci. 9,1154.

Tusher, V. G., Tibshirani, R., and Chu, G. (2001) Significance analysis of microarrays applied to the ionizing radiation response. Proc. Natl. Acad. Sci. U S A, 98(9), 5116–5121

Van der Weele, C.M., Spollen, W.G., Sharp, R.E. and Baskin, T.I. (2000) Growth of Arabidopsis thaliana seedlings under water deficit studied by control of water potential in nutrient-agar media. J Exp Bot. 51, 1555–1562

Verma, S., Gazara, R.K. and Verma, P.K. (2017) Transcription factor repertoire of necrotrophic fungal phytopathogen Ascochyta rabiei: predominance of MYB transcription factors as potential regulators of secretome. Front. Plant Sci. 8, 1037

Xia, J., Fjell, C. D., Mayer, M. L., Pena, O. M., Wishart, D. S., and Hancock, R. E. (2013). INMEX--a web-based tool for integrative meta-analysis of expression data. Nucleic Acids Res. 41, W63–W70.

Zheng, X., Yang, L., Li, Q., Ji, L., Tang, A., Zang, L., Deng, K., Zhou, J. and Zhang, Y. (2018) MIGS as a Simple and Efficient Method for Gene Silencing in Rice. Front. Plant Sci. 9:662.

Zhou, C.-M., Zhang, T.-Q., Wang, X., Yu, S., Lian, H., Tang, H., Feng, Z.-Y., Zozomova-Lihova, J. and Wang, J.-W. (2013) Molecular Basis of Age-Dependent Vernalization in Cardamine flexuosa. Science 340(6136), 1097–1100

